# Unsupervised Machine Learning for Adaptive Immune Receptors with immuneML

**DOI:** 10.64898/2026.04.15.718648

**Authors:** Milena Pavlović, Charlotte Würtzen, Chakravarthi Kanduri, Maria Mamica, Lonneke Scheffer, Christin Lund-Andersen, John Gubatan, Theresa Ullmann, Victor Greiff, Geir Kjetil Sandve

## Abstract

Machine learning (ML) enables adaptive immune receptor repertoires (AIRRs) analyses for biomarker identification and therapeutic development. With the majority of AIRR data partially or imperfectly labeled, unsupervised ML is essential for motif discovery, biologically meaningful clustering, and generation of novel receptor sequences. However, no unified framework for unsupervised ML exists in the AIRR field, hindering the assessment of model robustness and generalizability. Here, we present an immuneML release advancing unsupervised ML in the AIRR field through unified clustering workflows, interpretable generative modeling, integration with protein language model embeddings, dimensionality reduction, and visualization. We demonstrate immuneML’s utility in three use cases: (i) benchmarking generative models for epitope-specific sequence generation, assessing specificity and novelty, (ii) systematic evaluation of clustering approaches on experimental receptor sequences against biological properties, such as epitope specificity and MHC, and (iii) unsupervised analysis of an experimental AIRR dataset to examine potential confounding, a practice widespread in related fields but unexplored in AIRR analyses.

## Introduction

Adaptive immune receptors (AIRs) are proteins produced by B and T cells that recognize specific antigens (e.g., from viruses, bacteria, or cancer) and mount an immune response. A set of all AIRs in an individual constitutes an adaptive immune receptor repertoire (AIRR). It consists of ∼10^9^ unique AIRs^1–3^, indicative of both present and past immune responses. Understanding the rules that determine antigen binding and differentiating between AIRRs of individuals with and without certain diseases would open new opportunities for diagnostic, prognostic, and therapeutic applications^4^.

There are a few challenges to the AIRR analysis and understanding of these rules. First, observed AIR sequences reflect the combined influence of different biological factors: antigen recognition and presentation^5^ (e.g., via human leukocyte antigen (HLA)), V(D)J recombination process, cross-reactivity, high specificity, and vast diversity of the theoretical sequence space, and technical factors introduced during collection, sequencing and preprocessing of the data^6^. Second, while there is an increasing number of AIR(R) datasets deposited in publicly available databases such as VDJdb^7^, IEDB^8^, iReceptor^9^, the datasets are often only partially labeled (e.g., labels are available only on the repertoire level) or they might be imperfectly annotated^10^. These complexities, combined with the need to simultaneously consider multiple labels (e.g., HLA restriction, epitope specificity and batch effects), motivate the use of unsupervised approaches, such as clustering and representation learning, that can identify meaningful patterns and facilitate hypothesis generation. Furthermore, suitable approaches for generative modeling, typically trained in an unsupervised manner, may allow sampling of data with desired biological properties.

In the AIR(R) field, clustering has been used for several aims. For example, it is used to discover clusters of sequences with shared epitope specificity^11–16^. While specificity prediction is commonly formulated as a supervised learning task, clustering-based approaches remain relevant given the predominance of unlabeled AIR data, label noise, and receptor cross-reactivity, which can lead to multiple labels per receptor. Moreover, clustering may enable the identification of receptor groups associated with previously unseen epitopes. Beyond specificity prediction, clustering is used to identify receptors involved in ongoing immune responses^17^ and identify clonally related sequences^18^. While more common in other omics analyses^19^, clustering may also be used to detect batch effects in AIRR data. Clustering of repertoires is less common, but, in principle, it could be performed in the same way given suitable repertoire representations.

Defining AIR(R) representations is an open challenge. AIR sequence representations may be fixed, based on physicochemical features or sequence similarity^11,12,14,20^ (e.g., full sequence similarity, k-mer similarity, domain-inspired similarity metrics), or learned from the data. Learned representations can arise from deep learning models trained for specific tasks, or they can be derived in a self-supervised manner without reliance on predefined labels (e.g., from autoencoders or language models). Multiple such models have been proposed, including immune2vec^21^, ProtTrans^22^, ESM3^23^, AbLang^24^, AntiBERTa2^25^, SCEPTR^26^, SC-AIR-BERT^27^, TCR-VALID^28^, BertTCR^29^, TCR-BERT^30^, and GloVe-based representations^31^. While sequence similarity measures are traditionally used in AIR clustering applications, integrating learned embeddings into a common workflow to be used with classification or clustering approaches might be beneficial for AIR data analysis^32^.

Repertoire representations are usually either summary statistics of sequences in the repertoire^33,34^ or a subset of relevant sequences^35–37^. While repertoires are increasingly used for diagnostic purposes^4,38–43^, unsupervised repertoire analyses are less common.

Generative models have been employed to create new AIR(R) data. Some models follow the biological process of generating naive AIRs (OLGA^44^, IGoR^45^), the selection process (soNNia^46^), or the biological process followed by sequencing artefacts for benchmarking purposes^47^. Other models rely on variational autoencoders^48^ or other deep learning models for generating immune repertoires^49^ or epitope-specific receptor sequences^50–53^.

Despite recent advances in embeddings and algorithms, evaluating unsupervised approaches for both clustering and sequence generation remains challenging. Model selection for clustering and validation of clustering results is inherently difficult due to the lack of external supervision and many possible definitions of “true clusters”^54^. In bioinformatics applications, clustering often relies on single datasets and potentially misleading metrics^55–57^, without established processes for repeated runs or stability analyses. Furthermore, the evaluation of generative models is hindered by incomplete knowledge of biological rules, high dimensionality of the data, limited size of available datasets compared to the possible sequence space, high cost of experimental validation, and lack of standardized evaluation metrics capable of capturing different aspects of the generated sequences.

Collectively, these challenges underscore the need for systematic frameworks that enable comparison of unsupervised ML models across diverse feature representations, support robust exploratory analyses, incorporate stability assessment and validation of both results and model choice, provide a multitude of evaluation criteria, and facilitate seamless integration with downstream ML approaches.

To address these challenges, we introduce a unified framework for unsupervised ML analysis in the AIR(R) settings, integrated with and expanding the immuneML platform that was hitherto limited to (a broad range of) supervised ML analyses. To provide better insights into the generalizability and robustness of clustering results, the new release of the immuneML platform implements model selection for clustering based on stability assessment^58^ and resampling-based estimates of internal and external validation indices^59^. To validate clustering on a separate dataset, immuneML additionally implements the clustering validation framework formalized by Ullmann and colleagues^60^. To improve the interpretability of AIR generative models, it provides a generative modeling workflow along with visualization tools for comparing original and generated receptor sequences across multiple domain-specific criteria. The platform also integrates existing AIR clustering approaches, generative models, and embeddings based on protein language models (PLMs) into a unified framework, enabling reproducible and transparent analyses. Finally, we demonstrate the applicability of these workflows in three use cases: (i) benchmarking three generative models on simulated epitope-specific AIR sequences with known ground-truth signals determining the epitope specificity, (ii) exploring how experimental AIR sequences cluster with respect to known biological properties with simulated datasets providing a baseline, and, finally, (iii) employing unsupervised ML to investigate potential confounders in an experimental AIRR dataset.

## Results

immuneML is an open-source software platform for machine learning analysis of AIR(R) data^62^, with a strong focus on transparency and reproducibility of analyses. The original version supported multiple input formats, preprocessing, domain-specific encodings, and a classification workflow with model selection and assessment via a nested cross-validation scheme. The main applications were repertoire and receptor sequence classification (e.g., for disease diagnostics). This platform release extends these options to clustering workflows (e.g., for patient stratification, sequence specificity prediction, confounder analysis), generative models (e.g., for design of epitope-specific receptor sequences), embeddings based on protein language models (PLMs), and dimensionality reduction for data visualization and exploratory analysis (Figure 1a).

**Figure 1.**
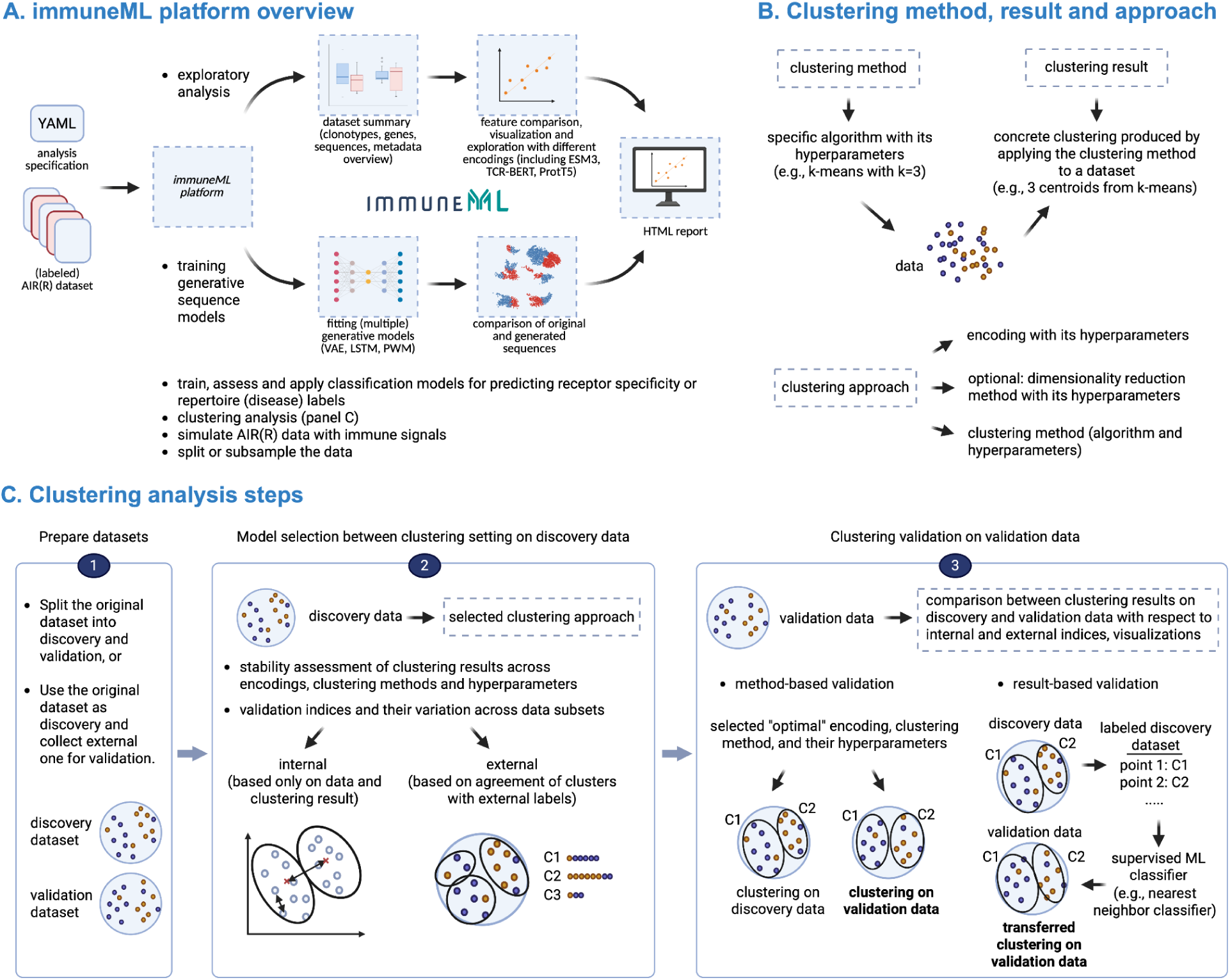
Overview of new immuneML features for dataset exploration and unsupervised machine learning analysis. **A**. The platform takes AIR(R) dataset and an analysis specification as input, performs the analysis and produces an HTML report as output. New features supporting unsupervised ML analyses include training generative sequence models, clustering analyses, LIgO integration^61^ for simulation of AIR(R) datasets, extended reporting for exploratory analyses, dimensionality reduction, new embeddings based on protein-language models. **B**. Focusing on clustering analysis, we illustrate the difference between clustering method, clustering result, and clustering approach. **C**. The clustering workflow in immuneML consists of three steps: preparation of separate discovery and validation datasets, model selection on discovery data and clustering validation on validation data. Model selection in the second step is based on the stability of the clustering^58^, validation indices across discovery data subsets^59^ and visualizations. Validation of the clustering is performed on separate validation data following the framework outlined by Ullmann and colleagues^60^, offering result-based and method-based validation.

Generative models of adaptive immune receptor sequences focus on sequence simulation for benchmarking purposes or designing sequences with desired properties, like antigen specificity or developability parameters^63,64^. Some such models are already available as independent tools^52,53^. However, it is important to examine these models and generated sequences to evaluate the learned models and to provide potential biological insights into the analyzed sequence sets. For that purpose, we introduce a workflow for generative models that can be easily combined with various data visualizations and statistical analysis reports. Further, we integrated the following models into our platform: SoNNia^46^, LSTM^50,51^, VAE^48^, ProGen2^65^. Additionally, the LIgO tool for AIR(R) simulation, which allows flexible simulation of both immune receptors and repertoires with known and fully annotated ground truth signals^61^ is integrated into the new immuneML version.

Building on the ability of generative models to design AIRs, the internal representation of AIRs of these models might be useful for capturing the underlying structural and functional properties and can be used in downstream analyses, such as classification or visualizations^32^. For that purpose, immuneML integrates several data representation techniques based on protein and AIR-language models: ProtT5^22^, TCR-Bert^30^, and ESM3^23^ are added to the Word2Vec-based encoding available in the original version.

Furthermore, immuneML now features dimensionality reduction techniques such as PCA, t-SNE^66^, and UMAP^67^, which can be combined with various data representation methods to examine the properties of the analyzed dataset (e.g., for interpreting the results of generative models or clustering approaches).

For clustering analysis of sequences and repertoires (Figure 1c), the new immuneML version provides a workflow for model selection across clustering approaches, which includes encoding, dimensionality reduction method, and clustering method, along with their hyperparameters (Figure 1b). The model selection is based on stability assessment as proposed by Lange and colleagues^58^: for many repetitions, the data is split into two equal parts, the clustering approach is fitted to both halves and then transferred in a supervised manner from one of them to the other. The correspondence between the two sets of labels is computed and the variability of it across repetitions is reported as the stability measure. In the immuneML implementation, this correspondence between two sets of labels is measured using adjusted Rand index^68^. The model selection procedure further includes data visualizations, and reporting of internal and external validation indices (measuring quality of the clustering based on the data alone, and measuring the agreement between cluster labels and external labels, respectively) computed over multiple data subsets^59^. Based on this information, the user selects the clustering approach. The dataset used for this purpose is referred to as discovery dataset. For model validation on the validation dataset, immuneML implements the framework proposed by Ullmann and colleagues^60^. The framework supports two types of validation: method-based and result-based. In method-based validation, clustering is performed independently on the discovery and validation datasets, and the results obtained for these two sets are directly compared. In result-based validation, a supervised classifier is fit to clusters determined on the discovery set and used to predict clustering on the validation set, indicating whether the clustering result is useful for the validation data. These two validation strategies provide complementary insight into the robustness and applicability of the chosen approach.

In many omics disciplines, exploratory unsupervised approaches, such as clustering or principal component analysis, are routinely applied to explore the hidden structure of the data and examine the influence of batch effects, protocol differences, or population stratification that may act as confounders for the analysis^19^. Comparable analyses have not yet become standard in AIRR research, despite a clear need to investigate the potential sources of bias in downstream applications^69^. We argue that exploratory and clustering analyses provided in immuneML represent a starting point for such investigations in the AIRR domain.

In addition to supporting unsupervised functionality, the new immuneML version significantly enhances the overall platform functionality available for both supervised and unsupervised analyses. Its new data model (Supplementary Figure 1) and data manipulation components internally rely on BioNumPy, an open-source Python package for efficient array programming on biological data^70^. This results in more extensive data validation and faster analysis. The usability and platform documentation have also been improved compared to the original version, through simplified installation options, new domain-specific data visualizations, tutorials, and examples.

### Use case 1: Comparison of generative models of antigen-specific TCR sequences

Simulated data with known ground-truth properties provides a principled basis for benchmarking of statistical and ML methods^71,72^. Building on this premise, we demonstrate the application of immuneML for generative modeling in the AIR domain by training and evaluating three distinct generative models on a simulated dataset with known ground-truth signals determining the antigen specificity (Figure 2a). The dataset was simulated using the LIgO^61^ simulation tool and included 5 distinct k-mer based signals of increasing complexity (see **Methods**). The generative models used were position-weight matrix (PWM), variational autoencoder (VAE) similar to Davidsen and colleagues^48^, and long short-term memory (LSTM) inspired by Saka, and Akbar and colleagues^50,51^. To assess the patterns the models learned, we generated 6000 sequences from each model and assessed the percentage of sequences containing the signals present in the original dataset, as well as whether the generated sequences containing those signals were memorized from the training set or unseen during training (novel) (Figure 2b). Among the datasets generated by the models, the LSTM dataset contained 98.88% of signal-specific sequences, evenly spread across the different signals. The VAE dataset contained 74.52% of signal-specific sequences, while PWM contained only 27%. However, while the LSTM dataset contained the highest percentage of signal-specific sequences (∼99%), roughly half of them were memorized (∼43%). In contrast, the majority of signal-specific sequences in the VAE dataset were not present in the original dataset (∼73%). The confusion matrix of a classifier trained to distinguish between sequences of different origins and UMAP visualization of 4-mer frequencies for the generated datasets additionally illustrates that while the original test dataset and LSTM dataset appear to overlap well, the VAE and, especially, the PWM datasets fall outside the original sequences (Figure 2c,d).

**Figure 2.**
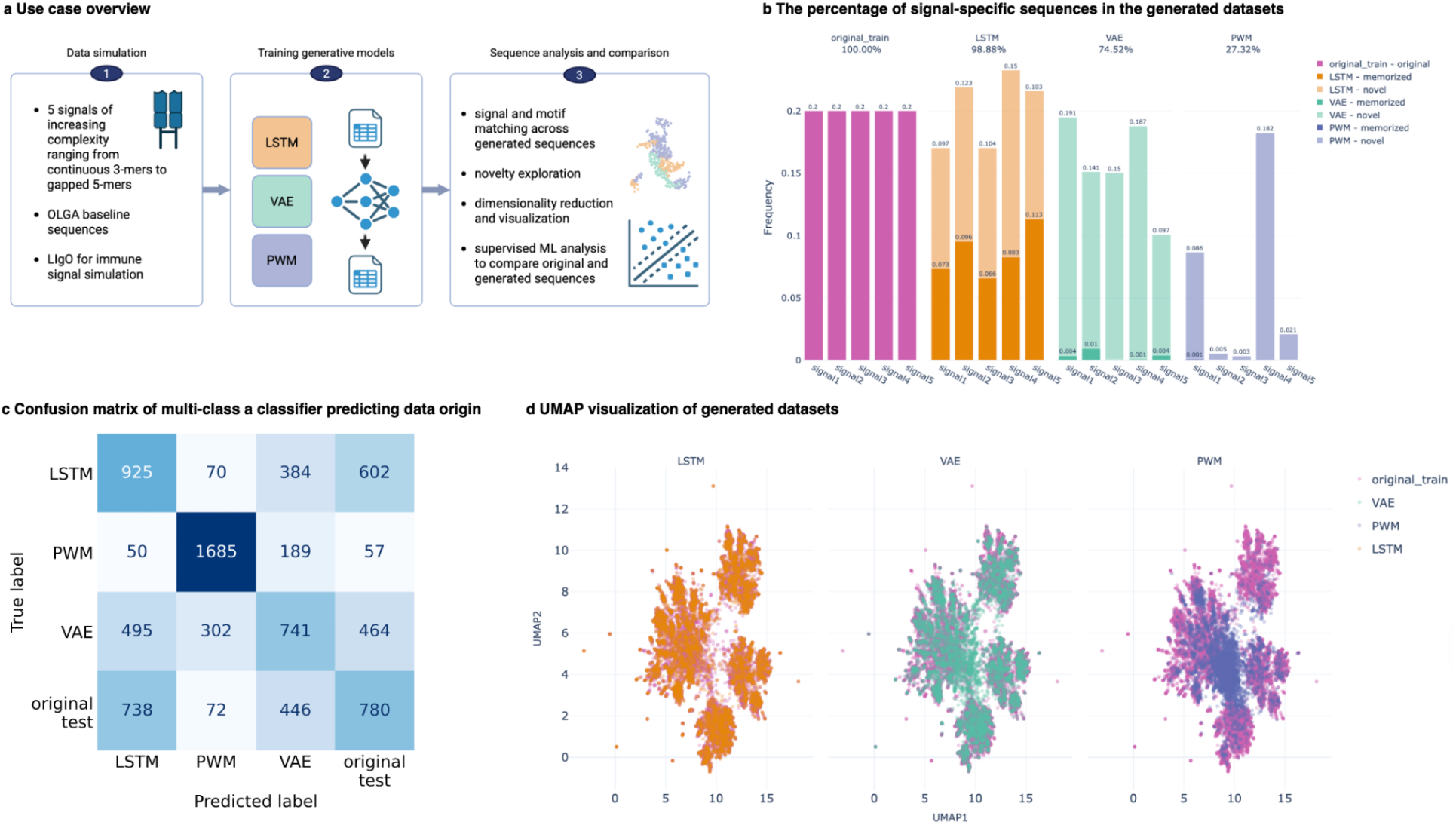
Analysis of the sequences produced by different generative models in use case 1. **a.** The use case overview: from simulating signal-specific sequences using LIgO with 5 different k-mers, to training multiple generative models, to analyzing the generated sequences and comparing them to the original sequences. **b**. The percentage of signal-specific sequences in the datasets generated by LSTM, VAE, and PWM models out of 6000 generated sequences. The original training dataset with balanced signals is shown for reference. The plot also highlights the fraction of novel sequences vs sequences memorized from the training in the generated datasets. **C**. The confusion matrix of multi-class logistic regression classifier based on 4-mer frequencies shows that it is possible to distinguish between the sequences of different origin, with PWM-generated sequences standing out the most. **d**. UMAP visualization based on 4-mer frequencies of the datasets generated by LSTM, VAE, and PWM models, with the original train dataset shown as a reference. The original dataset and LSTM dataset overlap almost completely, while the VAE covers some areas not present in the original dataset. PWM overlaps the least, consistent with its having the lowest percentage of signal-specific sequences.

### Use case 2: Exploring biological structure of epitope-specific TCRβ sequences using clustering

In the second use case, we evaluated the extent to which clustering captures biologically relevant information, such as epitope specificity or HLA restriction, using a publicly available dataset of TCRβ sequences from IEDB^8^. To do this, we explored multiple clustering approaches (including data encoding, optional dimensionality reduction technique, and clustering method along with their hyperparameters) across multiple data subsets to get insight into the robustness of the obtained results using immuneML’s clustering workflow (Figure 1c). We compared TCR-Bert, ProtT5, ESMC, kmer frequency-based encodings with PCA and k-means and tcrdist-based representation with hierarchical clustering with different numbers of clusters. We established a baseline performance of different approaches on four simulated datasets where we knew the characteristics guiding the clustering, and then applied it to the experimental IEDB dataset. We validated the findings on a separate experimental dataset also obtained from IEDB.

In simulated dataset 1 (random sequences of amino acids, see **Methods**) and simulated dataset 2 (OLGA-simulated TCRβ sequences with randomly assigned labels), we showed there is no correlation between clusters and assigned labels (Supplementary Figure 2). Strong clusters in simulated datasets 3 and 4 were successfully captured by the clustering approaches with highest median adjusted mutual information^73^ (AMI) of ∼0.8 (simulated dataset 4) and ∼0.5 (simulated dataset 3), both for tcrdist-based encoding with hierarchical clustering. Stability analyses on simulated datasets 1 and 2 showed that k-mer frequency with k-means and tcrdist with hierarchical clustering had low stability in absence of any signal. TCR-Bert, ProtT5, and ESMC encodings with k-means indicated increasing stability with lower number of clusters (median adjusted Rand index goes up to ∼0.5 for k=15 clusters). On simulated dataset 2 with OLGA-simulated sequences, this trend was similarly present, however, tcrdist approach exhibited higher variability, but it was still less stable than PLM-related approaches. On simulated dataset 4, stability as measured by adjusted Rand index went up to ∼0.9 for the tcrdist-based approach with 100 clusters. PLM-related approaches kept the same stability as in datasets 1 and 2, while tcrdist and k-mer frequency approaches became more stable for some numbers of clusters. The stability observed with PLM-related encodings may come from the properties of the embedding space (explored in more details elsewhere^74^) which produce a consistent structure on the data, partially independent of the data itself, especially evident from the analysis of simulated dataset 1 with random sequences (Supplementary Figure 3).

In the IEDB dataset, out of the explored clustering approaches, tcrdist-based encoding with hierarchical clustering best captured the epitope and HLA information, across different cluster sizes. However, AMI even for this clustering approach was only ∼0.14 (Figure 3b), which is much smaller than AMI in simulated datasets 3 and 4 with known, strong clusters (∼0.5 and

**Figure 3-.**
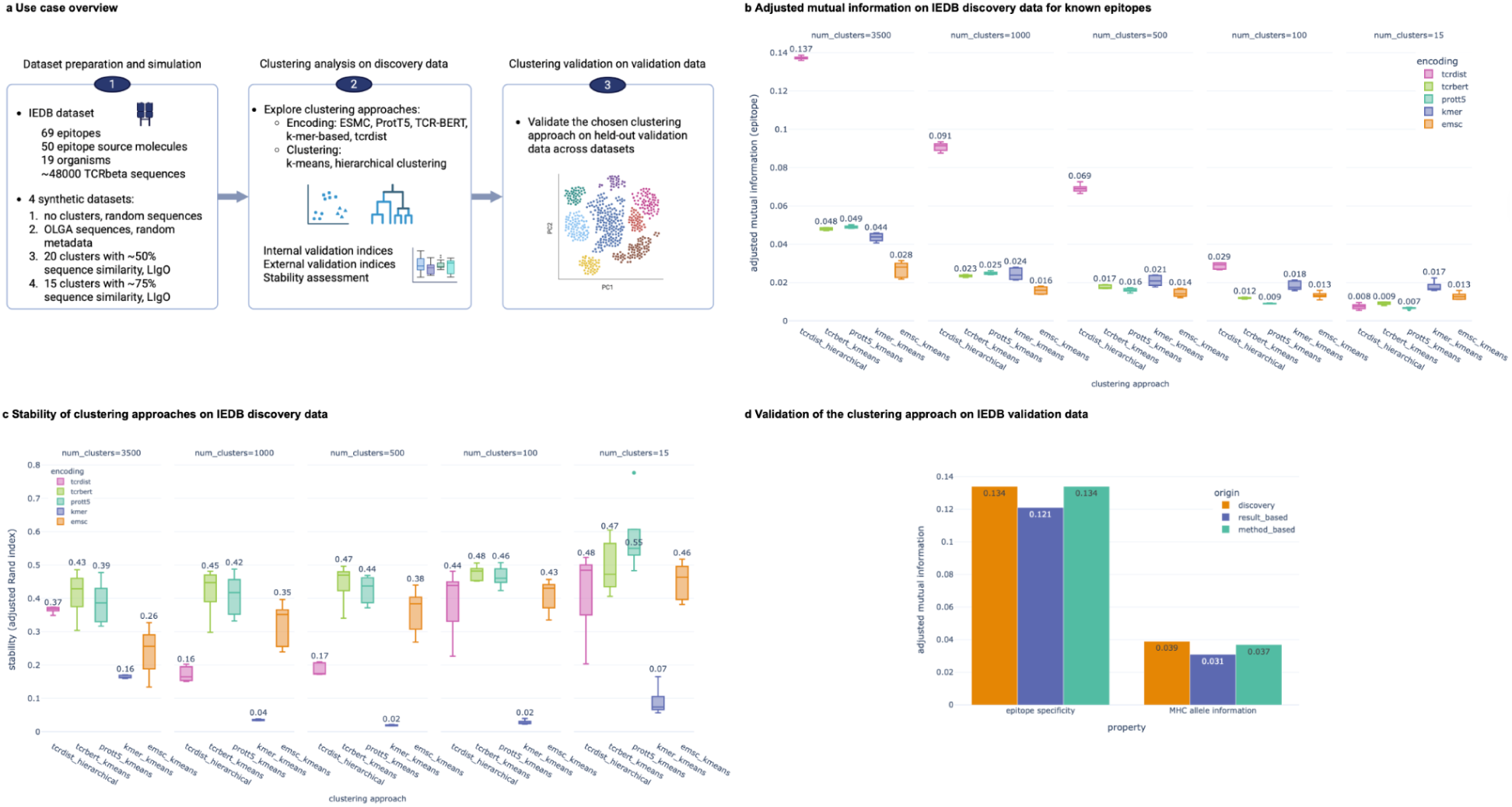
Clustering results on the IEDB dataset in use case 2. **a.** Overview of the use case: one experimental dataset for IEDB and four synthetic datasets were analyzed in the same workflow. The dataset was first split into discovery and validation, then clustering analysis was performed on discovery data where one clustering approach was chosen for validation on the validation (held-out) data. **b**. Adjusted mutual information (AMI) with respect to the epitope labels was computed across discovery data subsets, encodings, clustering methods and number of clusters, where the combination of tcrdist for data representation and hierarchical clustering consistently captured most of the epitope information. **c**. Stability of different encodings and clustering methods on IEDB discovery data. **d**. Selected clustering approach (tcrdist-based encoding with hierarchical clustering with 3500 clusters) performs similarly on discovery data and on held-out IEDB dataset for both method-based and result-based validation approaches, as measured by AMI with respect to biological properties (epitope specificity and MHC allele information).

∼0.8, respectively; Supplementary Figure 1). In terms of stability, clustering approaches based on PLM-based embeddings and k-means were the most stable, with tcr-dist-based encoding with hierarchical clustering being comparable, while k-mer encoding with k-means was the least stable (Figure 3c). Due to best correspondence to the expected main biological driver of clustering (epitope specificity) and similar performance for other, related metadata information (Supplementary Figure 4) and reasonable stability, we selected tcrdist-based encoding with hierarchical clustering (with 3500 clusters) for validation on an independent, held-out IEDB dataset. The selected approach for two labels of interest (epitope specificity and MHC allele information) performed similarly on discovery data and validation data for both method-based and result-based validation (Figure 3d).

### Use case 3: Analyses of confounders in an experimental AIRR dataset

In this use case, we demonstrated how immuneML supports exploration of confounding biases on experimental datasets. The focus of this use case is exclusively on initial explorations of potential undesired drivers of similarity, not considering any disease or treatment-related signals.

The dataset consists of paired single-cell TCR and BCR data from peripheral blood mononuclear cells (PBMCs) of inflammatory bowel disease (IBD) patients (n=143, with Crohn’s disease (CD) n=76, and ulcerative colitis (UC) n=67) and healthy controls (n=23). The dataset includes the information on the disease status, age, sex, sequencing batch and other properties. While the primary purpose of collecting this dataset was to investigate its potential utility for disease diagnostics, in the present work, it was instead used to demonstrate the initial exploratory analysis of an experimental dataset and to assess the influence of potential confounders prior to the application of supervised ML methods for diagnostics.

To show how the initial dataset analysis may be performed, we examined the distribution of the dataset’s external labels (disease status, age, sex, batch), their overlap, distributions of repertoire clonotype counts, gene usage, and explore dataset visualizations and feature comparisons for the BCR subset of the data (Figure 4a). From this analysis, we showed that some batches were exclusive to a single diagnosis status (Figure 4d) and that some of the batches clearly stood out (Figure 4b). To examine if batches leave such a strong imprint on the repertoires as to dominate overall similarity between repertoires, we performed clustering and compared the clustering results with three external labels using adjusted mutual information (Figure 4e) indicating that batch effects were indeed present across clustering approaches. However, upon examining the stability of clustering across multiple splits using a procedure proposed by Lange and colleagues and measured by adjusted Rand index to control for chance (Figure 4c), all clustering results were very unstable, indicating low confidence in the clustering results. We further examined the correspondence between the clustering result and batch information for the most stable clustering approach (positional 3-mer frequencies with k-means with k=14), showing that none of the batches were located in a single cluster, and the batch which had majority of repertoires in a single cluster (S1B), also included only diseased repertoires, which could be the additional driver of the clustering results (Figure 4f). This analysis indicates that while a few processing batches are associated with particular diagnosis statuses, the repertoires do not appear to inherit batch-related imprints (sequence characteristics) that would clearly distinguish them from other repertoires in the clustering analysis. It is still recommended to either correct for batch or report results stratified by batch for any subsequent supervised analysis, but the absence of prominent batch-related sequence signals indicates that both supervised and unsupervised analyses would be meaningful to perform on the full set of repertoires across batches.

**Figure 4-.**
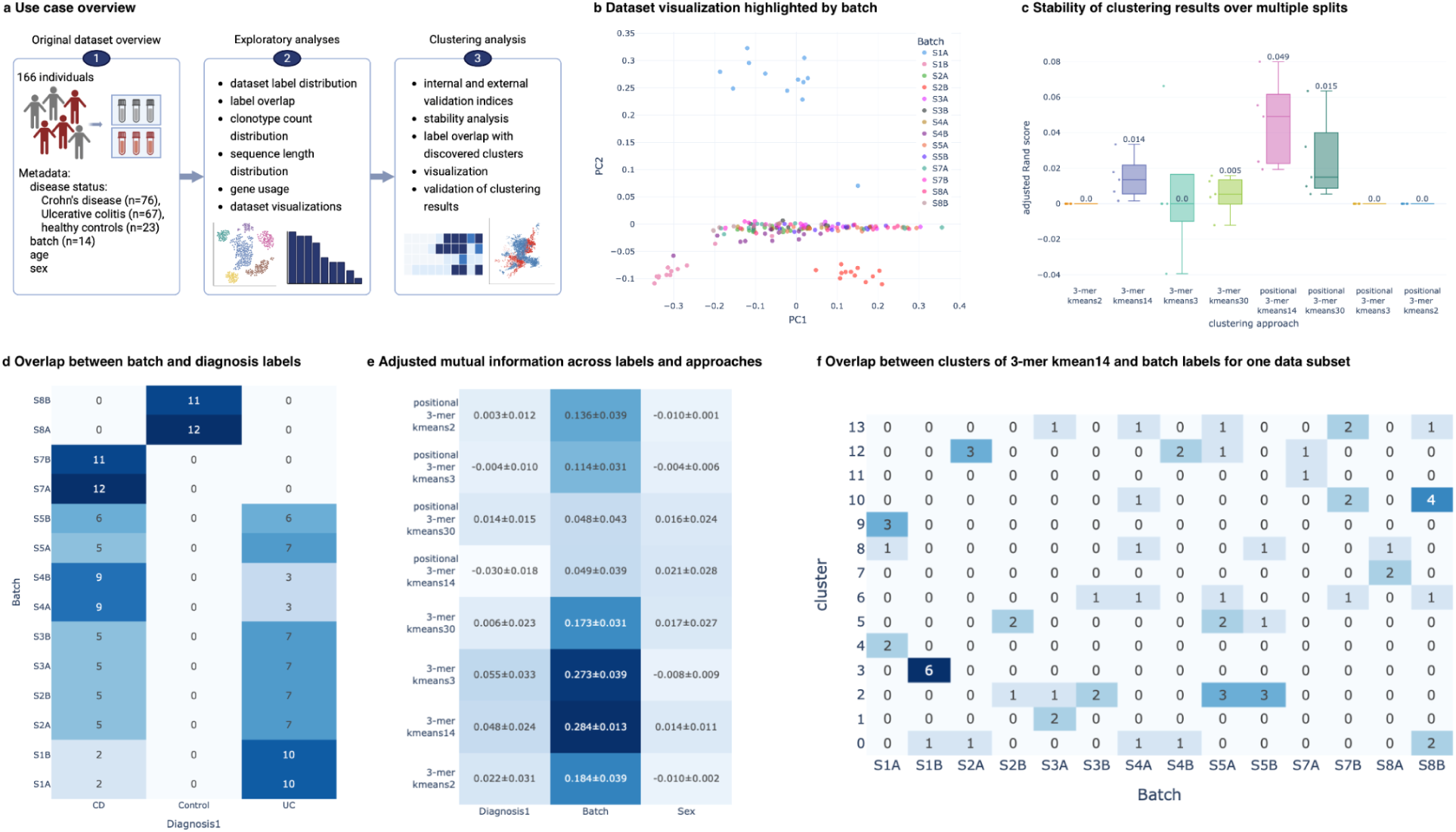
Confounder analysis of an experimental AIRR dataset. **a.** The dataset consists of 76 Crohn’s disease patients, 67 patients with ulcerative colitis and 23 healthy individuals. The workflow includes initial exploratory analysis, such as label distribution, label overlap, clonotype count distribution, sequence length distribution, gene usage, dataset visualizations and feature comparisons. If necessary, a subsequent (clustering) analysis can be performed if any potential issues were identified in the exploratory analysis. **b**. Dataset visualization based on positional 3-mer frequencies across repertoires with non-linear principal component analysis colored by batch shows that some batches clearly stand out (S1A, S1B, S2B). None of these batches contain healthy controls. **c**. Clustering is performed to assess if the batch effects dominate the signal. However, the stability analysis shows that clustering is very unstable. **d**. Overlap between batch and disease status (Diagnosis1) labels shows that some batches contain exclusively healthy controls (S8A, S8B), exclusively diseased repertoires of either Crohn’s disease or ulcerative colitis, or repertoires with Crohn’s disease only (S7A, S7B). **e**. Adjusted mutual information measures the similarity between clustering labels across clustering approaches and different metadata labels and indicates that batch effects are consistently detected by all clustering approaches to a different extent. **f**. Overlap between cluster labels and batch labels for one of the data subsets for the setting that was most correlated with the batch label (3-mer frequencies with k-means with k=14) shows that while some clusters contain the data only from a single batch (e.g., cluster 3 contains only S1B), none of the batches are present in only one of the clusters.

## Discussion

Unsupervised ML is a standard approach in biological sequence analysis, where it is applied to tasks such as motif discovery, clustering of sequences or samples, dimensionality reduction for visualization or downstream analyses, and identification of potential biases. Similar applications are increasingly being explored in AIR(R) research. However, a standardized framework for unsupervised ML in AIR(R) analysis is lacking, and current practices do not consistently reflect established standards in unsupervised ML^75^.

Several AIR(R) analysis packages have been developed for exploratory analyses and selected unsupervised learning tasks^9,76–81^, focusing on preprocessing, summary statistics, diversity analysis, repertoire overlap, phylogenetic analysis, dimensionality reduction, visualization, and limited clustering analyses. While these tools provide valuable functionality, they do not support the systematic evaluation of unsupervised ML methods for AIR(R) data.

immuneML addresses this gap by providing rigorous, reproducible workflows that critically assess the generalizability of unsupervised learning results, in line with established best practices from statistics and ML^75^. The platform emphasizes transparency and interoperability, offering the first unified framework for unsupervised learning in AIR(R) analysis. The latest version expands its functionality to include unsupervised workflows for training generative models, performing clustering analyses, and improving data representation and visualization, as demonstrated by the three use cases. This unification not only streamlines workflows but also improves usability, ensures comparability across studies, and supports more robust and reproducible discoveries.

While immuneML provides a robust clustering workflow, some domain-specific clustering tools are not available on the platform due to restrictive licenses, implementation in other programming languages, lack of packaging, and dependence on older package versions. Tools such as tcrdist3 are included, and for others, compatibility is supported through common data formats. immuneML also integrates data representation options based on language models and generative models, with more models available outside the platform. Integration is straightforward, with detailed developer tutorials available in the documentation. We invite the community to contribute to the platform with new models.

immuneML represents a step toward systematic unsupervised analyses in AIR(R) research. Early efforts, such as a recent benchmarking study of clustering approaches^82^, illustrate the potential for structured evaluation, but even these are limited in scope due to reliance on a single experimental dataset or a small set of metrics. More generally, unsupervised ML is challenging as evaluation indices might be biased, for example, favouring solutions with more clusters or poorly matching the domain knowledge^56,57^. Some of the concerns about the use of evaluation indices are also raised in the AIRR field^83^. By supporting a variety of validation indices, clustering stability analyses, different validation strategies, and detailed reporting, immuneML facilitates addressing this challenge. For generative models, domain-informed representations, comparisons and reporting are critical to ensure biologically meaningful results, and immuneML contributes to this effort. Together, these capabilities allow immuneML to offer structured workflows for supervised, unsupervised, and generative model analyses, enabling rigorous and reproducible ML applications within the AIRR field.

## Methods

### immuneML availability

immuneML is available as a Python package on PyPI (https://pypi.org/project/immuneML/). The code is available on GitHub (https://github.com/uio-bmi/immuneml). immuneML is also available on DockerHub (http://registry.hub.docker.com/r/milenapavlovic/immuneml) and conda (https://anaconda.org/bioconda/immuneml).

### immuneML architecture

immuneML takes in a YAML specification as input and outputs an HTML overview of the results. In addition to the analysis results, immuneML outputs a specification file containing the resolved default parameters, enabling full reproducibility of the analysis.

The platform was developed with the focus on extendability and interoperability: it supports a variety of input data formats, most notably the community AIRR format^84^, and exports the generated data in the same format so that it can be used by other AIRR tools. All intermediate results, raw data behind the generated plots, predictions made by the models, and the models themselves are also exported.

immuneML’s internal data representation offers two options: an object-oriented view of the data and support for array programming based on the BioNumPy library^70^ (Supplementary Figure 1).

The analyses are organized into instructions that include: classifier selection and assessment workflow (TrainMLModel), application of trained classifiers to new datasets (ApplyMLModel), preprocessing, encoding and visualizations of datasets (ExploratoryAnalysis), training generative model with optional post-analysis (TrainGenModel), generation of sequences using an already trained model (ApplyGenModel), the clustering workflow described in **Results** (Clustering), a convenience instruction for constructing smaller datasets (Subsampling), simulation of AIR(R)s with known ground-truth immune signals (LigoSimulation), and an instruction to assess the feasibility of the simulation with LIgO (FeasibilitySummary).

Machine learning models, encodings, and reports in immuneML implement the corresponding abstract classes, which define the functions and arguments necessary for the new classes to work seamlessly with the rest of the platform. In existing implementations, immuneML relies heavily on general data processing, visualization, and ML libraries such as PyTorch^85^, scikit-learn^86^, transformers^87^, pandas^88^, NumPy^89^, scipy^90^, gensim^91^, and Plotly^92^.

To extend the platform with new ML methods, data encodings, and reports, we provide detailed developer documentation (https://docs.immuneml.uio.no/latest/developer_docs.html).

### Use cases

#### Use case 1: Benchmarking generative models on ground-truth synthetic data

To demonstrate the utility of immuneML for training and evaluating generative models, and to highlight how the platform enables impartial and reproducible method comparisons, we benchmarked three different families of generative models on synthetic data with known ground-truth immune signals. The original sequence dataset was created using immuneML’s LIgO simulation instruction, employing OLGA^44^ to generate the background 20,000 human TCR beta amino acid sequences. Five immune signals of varying motif complexity were incorporated during simulation through rejection sampling. The signals included both contiguous and gapped motif patterns, ranging from 3- to 5-mers: *DEQ, GQET, NQPQH, AS–G,* and *SS-GT*. Each motif was found in 4,000 receptor sequences, resulting in equal representation of all signals. The dataset was split to preserve signal proportions between train and test sets, where 70% of sequences were used for training.

For benchmarking, we fitted three representative generative methods: a Long Short-Term Memory (LSTM) model, a Variational Autoencoder (VAE), and a Position Weight Matrix (PWM) as a statistical baseline. From each trained model, we sampled 6,000 sequences and assessed model performance by comparing generated distributions against both train and test distributions using domain-specific exploratory analyses.

To evaluate how well the models capture the respective immune signals of different complexities, we used immuneML’s built-in true motifs summary barplot report, which quantifies the frequency of generated sequences containing the ground-truth motifs. This analysis further distinguishes between *memorized* sequences (exact matches to training data) and *novel* sequences (previously unseen in training).

Additional analyses compared the feature distributions derived from different k-mer encodings, providing complementary perspectives on how well each model reproduced relevant sequence properties across signal types.

#### Use case 2: Exploring biological structure of epitope-specific TCRβ sequences using clustering

To demonstrate the utility of immuneML’s clustering workflow, we performed cluster validation analysis on both experimental and synthetic data. Below, we describe the datasets, clustering approaches, validation indices, and experiments performed.

##### Datasets

We perform the analysis on five datasets, one experimental and four simulated:

- IEDB dataset: ∼48000 human TCRβ sequences downloaded from the IEDB database (February 2nd, 2026) binding to 69 different epitopes, sourced from 50 different antigens from 19 different organisms.
- Simulated dataset 1: 10000 random amino acid sequences with no biological interpretation of length 14 or 15, with 20 different randomly-assigned labels with uniform probability.
- Simulated dataset 2: ∼24000 TCRβ sequences from the OLGA model with randomly assigned labels that correspond to metadata available in the IEDB dataset.
- Simulated dataset 3: ∼22000 TCRβ CDR3 sequences with 20 clusters of approximately equal size, where each cluster is enriched for one of the 20 randomly selected 4-mers. On average, approximately ∼50% sequence similarity is observed within each cluster. The dataset is simulated using LIgO tool available in immuneML.
- Simulated dataset 4: ∼15000 TCRβ CDR3 sequences with 15 strong clusters of approximately equal size, where each cluster is enriched for one of the 15 pairs of 4-mers (i.e., each sequence contains 1 of the 15 pairs of 4-mers). On average, approximately 75% sequence similarity is observed within each cluster. The dataset is simulated using LIgO tool available in immuneML.

##### Clustering approaches

We used multiple combinations of encodings, clustering methods, and a predefined number of clusters, resulting in a large number of clustering approaches. The immune receptor sequences were encoded into numerical vectors or pre-computed distances using a variety of approaches, including different protein language models (Prot-5^22^, ESMC^93^), fine-tuned language models for TCRs (TCR-BERT^30^), k-mer frequencies, and pre-computed distances using tcrdist3^13^. Except for pre-computed distances, the encoded data is reduced to fewer dimensions using PCA. k-means and complete-linkage agglomerative clustering have been used to cluster the encoded data into a pre-specified number of clusters (n = 15, 100, 500, 1000, 3500).

##### Validation indices

To assess the quality of clustering results, we used external validation indices to measure how well the clusters capture the available biological or external information. For this purpose, we used adjusted mutual information score (AMI), which measures the amount of information shared between two clusterings^73^. AMI is adjusted for chance and can thus be safely used for cluster validation analysis. For stability assessment, we used adjusted Rand index^68^. To measure the quality of the clustering results, we additionally report Silhouette and Calinski-Harabasz scores.

##### Analyses

The analyses are performed in three steps for each of the datasets: (i) splitting the dataset into discovery and validation subsets, with 50% split between the subsets, (ii) running clustering on the discovery data for all clustering approaches and reporting the full results across external metadata available for the dataset, internal indices and stability analyses, and (iii) validating the clustering by applying a selected clustering approach to the validation data (method-based validation) and transferring the clusters from discovery data to validation data (result-based validation).

#### Use case 3: Analyses of confounders in an experimental AIRR dataset

##### Dataset

The dataset consists of paired single-cell TCR and BCR data from peripheral blood mononuclear cells (PBMCs) of inflammatory bowel disease (IBD) patients (n=143, with Crohn’s disease (CD) n=76, and ulcerative colitis (UC) n=67) and healthy controls (n=23) in AIRR data format. For the subsequent analyses, we focus on BCR data.

##### Clustering approaches

To encode the data, we used positional and non-positional 3-mer frequencies. For dimensionality reduction, we used PCA with radial basis function kernel. For clustering methods, we used k-means where k was varied to correspond to the expected number of clusters (2 if clustering by sex, 3 if by disease status, 14 if by batch, or 30 in case a finer-grained clustering was beneficial).

##### Analyses

We conducted four different analyses of this dataset: (i) filtering the dataset to contain only BCRs as the repertoire files included a few of TCR sequences, (ii) exploratory analysis to examine the distribution of metadata labels, overlap of different labels (batch, diagnosis, sex), clonotype counts across disease statuses and batches, Shannon diversity, sequence length distribution by disease status and locus, gene usage distribution, (iii) exploratory analysis of encoded repertoires using positional 4-mer frequencies, non-positional 3-mer frequencies, in combination with non-linear PCA for dimensionality reduction for dataset visualization highlighted by different metadata labels of interest. Finally, seeing that there is a potential for confounding by the sequencing batch, we ran a clustering analysis in which we explored different encoding, dimensionality reduction and clustering methods to assess the degree to which the clustering is guided by the sequencing batch, as described under **Clustering approaches**. We performed clustering stability analyses and method-based and result-based validation as described in **Results** and by Ullmann and colleagues^60^.

## Data availability

The specification files and instructions how to reproduce the results for all three use cases are available in a dedicated GitHub repository (https://github.com/immuneML/immuneML-unsupervisedML-usecases). The data and results for the use cases are available on Zenodo at 10.5281/zenodo.19564623 (use case 1) and 10.5281/zenodo.19565450 and 10.5281/zenodo.19590967 (use case 2). The dataset used in use case 3 is a part of the manuscript in preparation and will be made public along with the results upon submission of that manuscript.

## Code availability

The immuneML source code is openly available on GitHub (github.com/uio-bmi/immuneML) under a free software license (AGPL-3.0). The immuneML Python package can be downloaded from pypi.org/project/immuneML.

## Supplementary figures

**Supplementary Figure 1 -.**
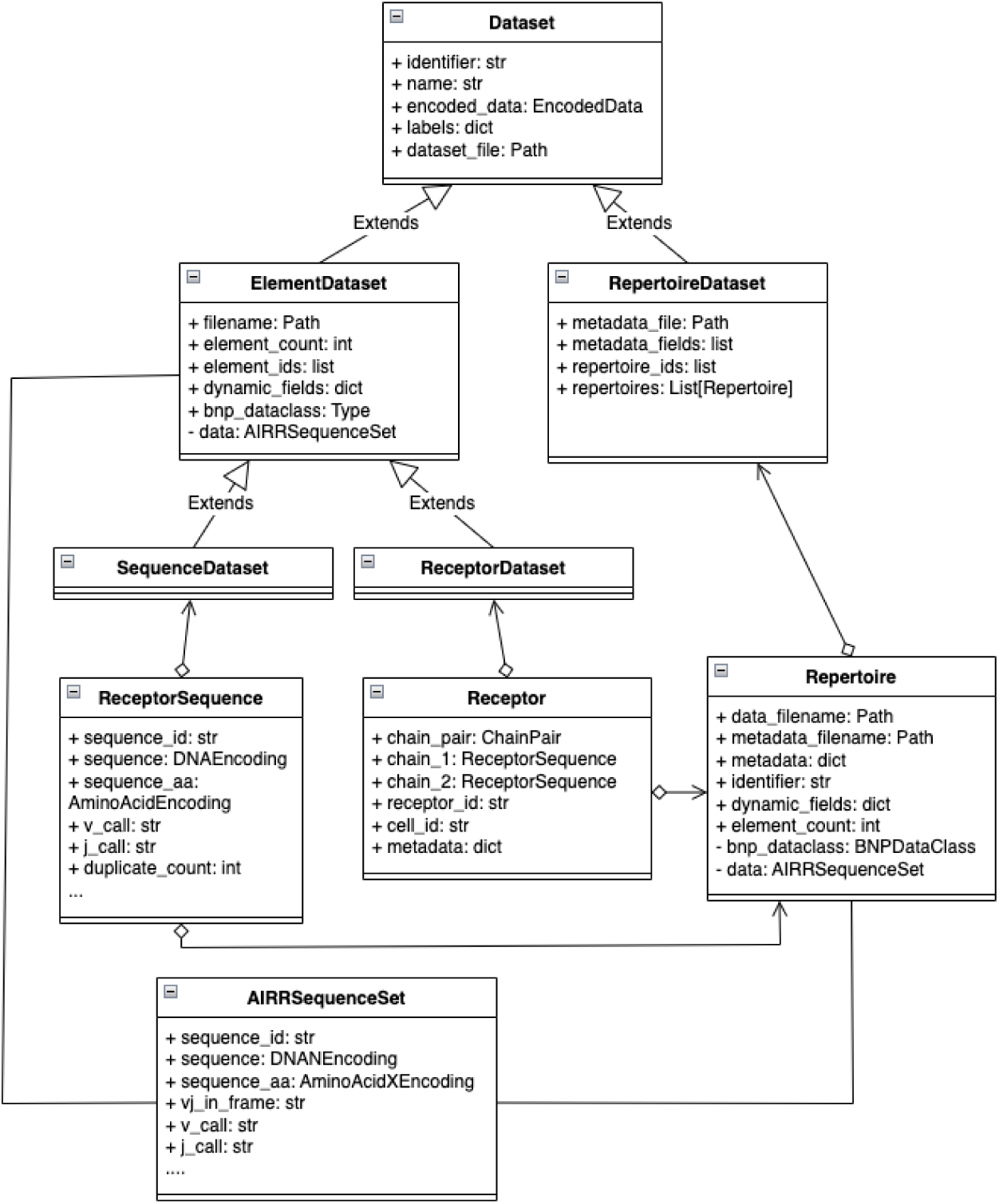
Data model class diagram. There are three dataset types: SequenceDataset, ReceptorDataset, and RepertoireDataset for different levels of analysis. AIRRSequenceSet is a data class based on bionumpy and is used by the SequenceDataset, ReceptorDataset, and the Repertoire class to access all the AIR sequences at once. ReceptorSequence and Receptor classes are utility classes for working on the object level; however, AIRRSequenceSet is available in the same places and can be used to speed up the analyses.

**Supplementary Figure 2 -.**
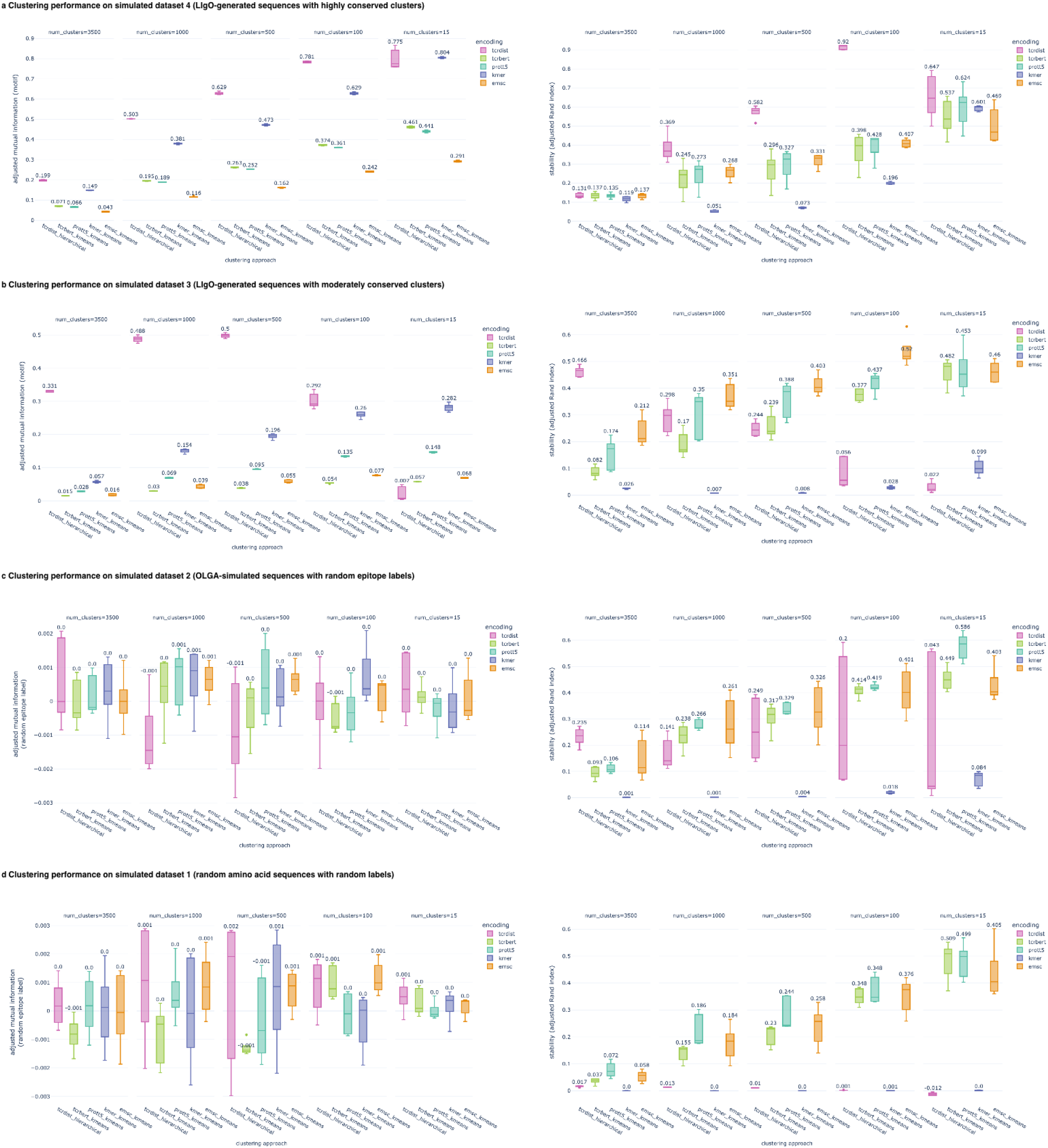
Clustering performance on simulated datasets from use case 2, as measured by adjusted mutual information (AMI) between a label of interest and the cluster labels obtained by a specific clustering approach, and stability (measured by adjusted Rand index (ARI)) across 5 repetitions. **a**. For simulated dataset 4, with 15 strong clusters, where the label of interest is simulated as a pair of 4-mers, AMI for all considered clustering approaches is higher when the number of clusters is closer to the ground truth. Stability is also higher when the number of clusters closer to 15. **b**. For simulated dataset 3, with 20 true clusters, where the label of interest is simulated as a single k-mer, k-mer frequency with k-means has increasing AMI as it approaches the number of true clusters. The same applies to all other clustering approaches, with the exception of tcrdist-based approach, which performs best overall, but only when the number of clusters is 500 or 1000. The stability of the clustering approaches is higher for fewer clusters, except for tcrdist-based approach which follows the opposite trend. **c**. For simulated dataset 2 with realistic OLGA-generated sequences and randomly assigned labels, AMI is close to 0 for all clustering approaches. Stability is higher for fewer clusters, but is less than stability with highly conserved clusters as shown in panel a. **d**. For simulated dataset 1 with random amino acid sequences and randomly assigned labels, AMI is close to 0 for all clustering approaches. Stability is higher with fewer number of clusters for clustering approaches based on protein language models, but close to 0 for k-mer frequency with k-means and for tcrdist-based approach.

**Supplementary Figure 3 -.**
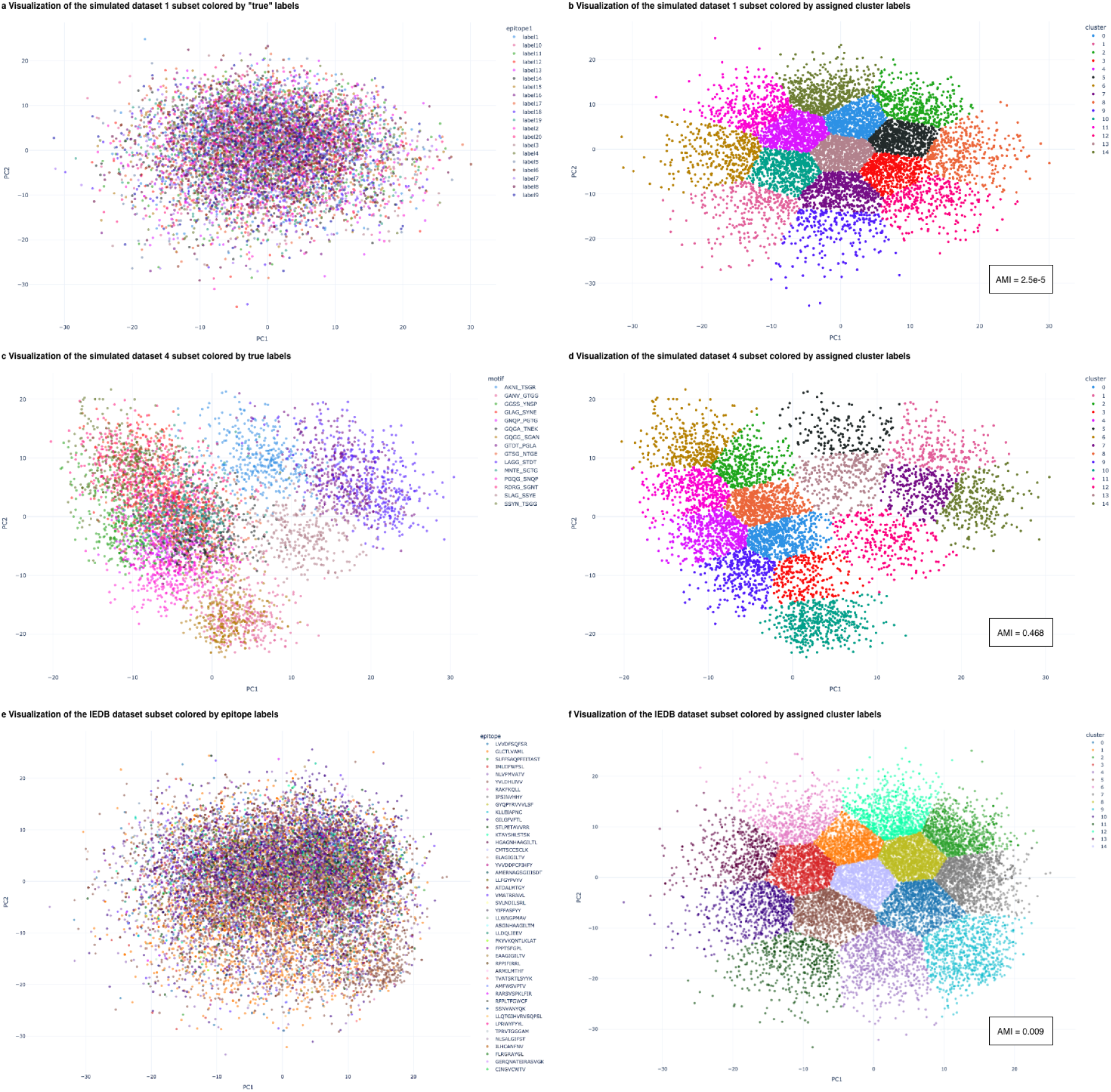
Visualization of clustering results and “true” labels across datasets provides insight into stability performance of protein language model-based encodings, shown here for TCR-Bert encoding, PCA with 2 components and k-means with k=15. **a**. Visualization of a subset of the simulated dataset 1 with random amino acid sequences and randomly assigned labels. **b.** Visualization of the same data as in panel a, colored by assigned cluster labels. As there is no structure in the data, the only structure present is the one imposed by encoding, which is results in similar clusterings by k-means and causes the high stability as seen in Supplementary Figure 2d. The adjusted mutual information (AMI) is 2.5*10^−5^. **c**. Visualization of a subset of the simulated dataset 4 with amino acid motifs present in realistic TCRβ sequences, with 15 true clusters. **d**. Visualization of the same dataset as in c, colored by assigned cluster labels, where the true clusters are successfully recovered (AMI=0.468). **e**. Visualization of a subset of the IEDB dataset colored by epitope labels. **f**. Visualization of the same dataset as in e, colored by assigned cluster labels. While relatively stable, this clustering does not correspond well to epitope labels (AMI=0.009).

**Supplementary Figure 4 -.**
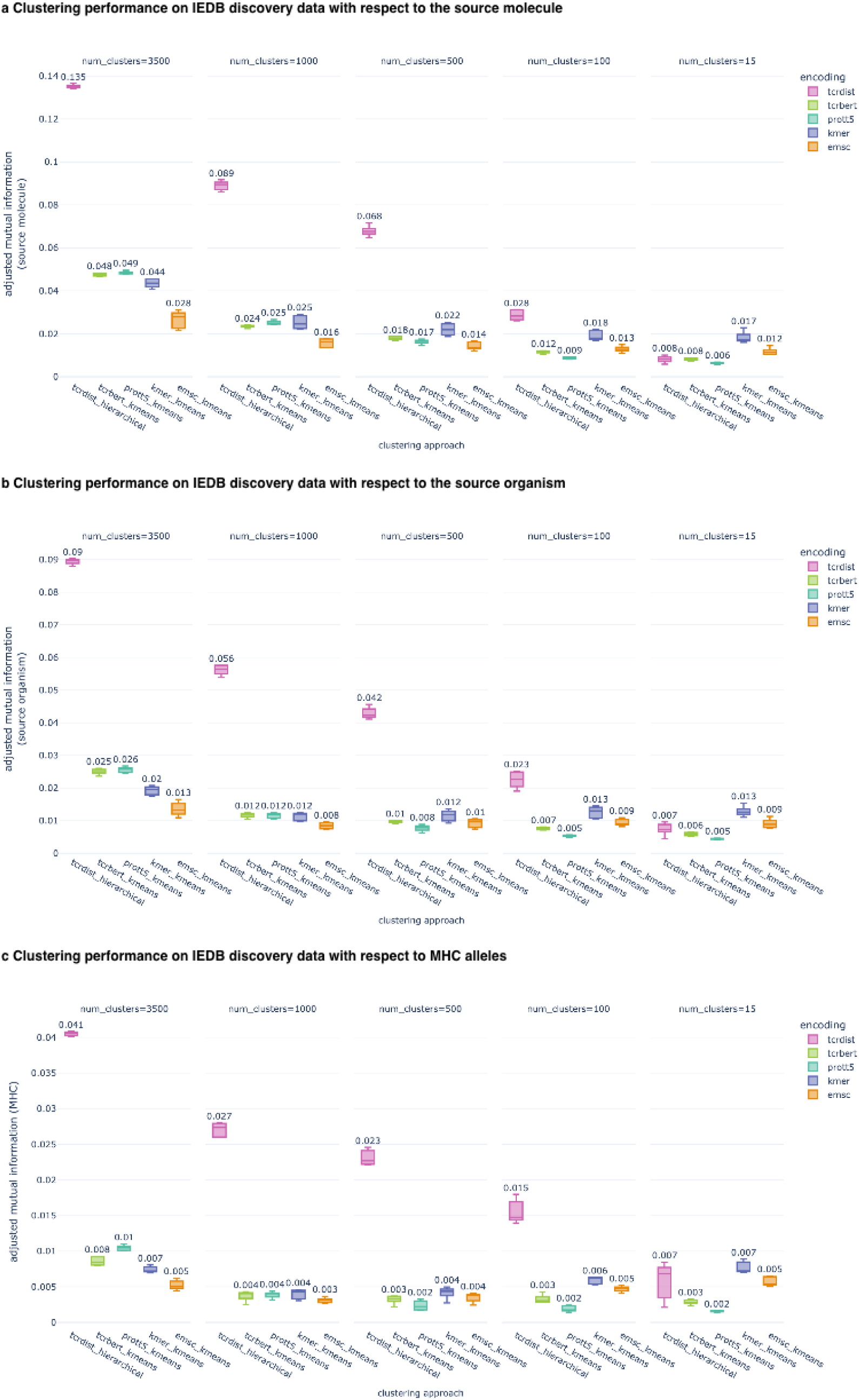
Clustering performance on IEDB data with respect to additional metadata labels indicates consistent performance by tcrdist-based approach across the labels. **a**. With respect to the epitope source molecule, the adjusted mutual information (AMI) of ∼0.135 is similar to performance with respect to epitope. **b**. With respect to the epitope source organism, AMI of ∼0.09 is lower than for epitope, but comparable. **c**. With respect to MHC alleles, tcrdist-based approach also captures the most information, with highest AMI ∼0.04.

